# The vaccinia virus E3L dsRNA binding protein detects distinct production patterns of exogenous and endogenous dsRNA

**DOI:** 10.1101/2023.09.21.557600

**Authors:** Wenjing Zhang, Nikhila S. Tanneti, Alejandra Fausto, Joangela Nouel, Hanako Reyes, Susan R. Weiss, Yize Li

## Abstract

Double-stranded RNA (dsRNA) is a pathogen associated molecular pattern recognized by multiple pattern recognition receptors and induces innate immune responses. Viral infections can generate dsRNA during virus replication. Genetic mutations can also lead to endogenous dsRNA accumulation. DsRNA is present in multiple conformations such as the A form (A-dsRNA) or Z form (Z-dsRNA). A-dsRNA has been detected from multiple viruses with positive-stranded RNA genomes (+ssRNA) but rarely from viruses with negative RNA genomes (-RNA); Z-dsRNA can be detected from influenza virus and poxvirus infections. Viruses have evolved mechanisms to antagonize cellular antiviral responses triggered by dsRNAs. For example, the vaccinia-virus E3L protein can bind and sequester dsRNA to evade host immune responses. The E3L protein encodes a Z-DNA and a dsRNA binding domains that bind to Z-form nucleic acids or dsRNA, respectively. Here we developed recombinant E3L proteins to detect dsRNA and Z-dsRNA generated from viral infections or endogenous cellular mutations. We demonstrate that the E3L recombinant protein specifically detects A-dsRNA generated from +ssRNA viruses but not-RNA viruses. We observe that among various virus infections assayed, only the influenza A virus generates Z-RNA that can be detected by anti-Z-NA antibody but not by the E3L recombinant protein containing the Z-DNA domain. The E3L recombinant protein can also detect endogenous dsRNA in PNPT1 or SUV3L1 knockout cells. Together we concluded that A-dsRNA can be produced and detected from viruses with +ssRNA genomes but not-RNA genomes, and Z-dsRNA can be produced and detected from influenza A virus.

**Importance:** The detection of dsRNAs, which exist in the A-dsRNA or Z-RNA conformation, is important for the induction of innate immune responses. dsRNA are generated during a virus infection due to virus replication, or can accumulate to genetic mutations. We engineered recombinant vaccinia virus E3L protein that can detect A-dsRNA generated during infection with a positive-sense RNA genome virus but not a negative-sense RNA genome virus. Infection with influenza A virus generates Z-RNA that can be detected with an anti-z-antibody but not the E3L recombinant protein. The E3L recombinant protein also detects endogenous dsRNA in PNPT1 or SUV3L knockout cells. These findings highlight important characteristics of dsRNA structure and detection.

## Introduction

The presence of double-stranded RNAs (dsRNAs) in cells is a hallmark of viral infections. Viruses composed of a dsRNA genome deposit long dsRNAs upon infection, while viruses with single-stranded genomes generate dsRNA as replication intermediates (1). Some DNA viruses can also generate dsRNA from overlapping transcripts and secondary structures during virus replication (2). Endogenous dsRNAs, while less abundant, can be found under certain cellular conditions or genetic mutations. For example, endogenous retroelements generate RNAs that serve as Zα-ligands and activate Z-DNA Binding Protein 1 (ZBP1) (3, 4). Genetic mutations can also lead to the accumulation of endogenous dsRNA; for example, mutations in SUV3L1 and PNPT1 lead to the accumulation of mitochondrial dsRNA that can escape into the cytoplasm leading to activation of MDA5-driven antiviral interferon responses (5). While mammalian cells readily recognize and tightly regulate the expression of DNA and short single-stranded RNAs generated during various cellular processes, they do not generate long dsRNAs. As dsRNA are typically generated during pathogenic conditions, cells have evolved mechanisms to detect dsRNA and trigger a cascade of antiviral responses to defend the host from pathogenic viruses. DsRNAs are present in either the A-form or Z-form (6). A-RNAs are recognized by signaling pathways that trigger antiviral responses, such as MDA5 RIG-I, OAS-RNase L, or PKR. ZBP1, a nuclear sensor of Z-NAs, recognizes Z-RNA during influenza A virus (IAV) replication and activates RIPK3-mediated nuclear rupture and necroptosis (7). The host ADAR p150 isoform recognizes the left-handed conformation of Z-RNA and prevents the interferon and innate immune responses mediated by MDA5 and ZBP1(8, 9). Viruses have evolved ways to antagonize host antiviral responses, for example, the vaccinia virus E3L protein sequesters dsRNA through dsRNA binding domain and Z-NA binding domain to evade antiviral activation of interferon, OAS, PKR and ZBP1 mediated pathways (10–12).

Currently, we have limited tools to detect dsRNA. The most commonly available marker of dsRNA is the J2 antibody, which can detect dsRNA from cells infected with many RNA or DNA viruses (13). Alternatively, the 9D5 antibody has been reported to have higher specificity to dsRNA compared to the traditional J2 antibody. Interestingly, while detection of dsRNA generated by negative-sense RNA virus infections is unsuccessful with J2 antibody, it is possible with the 9D5 antibody (14). However, the influenza A virus, a negative-sense RNA virus, generates Z-RNAs during replication that are not recognized by either J2 or 9D5. Exogenous viral proteins that bind to dsRNA can also be used to detect dsRNAs. The B2 protein from nidoviruses can detect dsRNAs with high sensitivity, but it is limited to dsRNA generated from positive-sense virus infections (1, 15). Thus, there is a need for probes that can detect dsRNAs from positive-sense RNA virus genomes. Such probes can also help us understand structural differences between positive- and negative-sense RNAs, which can help us understand how they trigger and surmount different immunological responses in the cell. Here, we generated an array of recombinant proteins expressing the vaccinia virus E3L protein and its derivatives that include the E3L dsRNA-binding motif, dsDNA-binding motif, and its Z-DNA-binding motif. These recombinant proteins were assayed for their ability to help visualize various populations of RNA in mammalian cells. We propose the recombinant E3L protein as a novel tool to detect A-dsRNAs and but not Z-dsRNAs generated during infections by RNA viruses and endogenous processes such as mitochondrial mutations. The study also raises important questions about the differences in RNA biology between positive and negative sense RNA viruses.

## Results

### Generation of recombinant E3L proteins

To assess the ability of the vaccinia virus E3L protein to bind and detect dsRNA, we generated multiple recombinant E3L proteins **(Figure 1A)**. All recombinant proteins contain the Maltose-binding protein (MBP) tag at the N-terminal to facilitate detection of the recombinant proteins with anti-MBP or anti-V5 antibodies. The E3L recombinant protein contains the full-length E3L protein from vaccinia virus which includes the Z-DNA binding domain and dsRNA binding motif. The E3L-mutant recombinant protein contains K167A and R168A mutations that inactivate the dsRNA binding motif or E3L (12). The E3L-zDNA protein contains only the Z-DNA binding domain of the protein. The E3L-dsRNA protein contains the dsRNA binding domain of the E3L protein. The B2 protein contains the dsRNA binding domain from the Flock House virus (1, 15). All recombinant proteins contain the Maltose-binding protein (MBP) tag at the N-terminal and a V5 tag at the C-terminal, which facilitates detection of the recombinant proteins with an anti-MBP antibody or the V5 antibody. All recombinant proteins were generated in E. coli and purified **(Figure 1B)** per protocol. The purified proteins were used in immunofluorescence protocol and visualized by anti-MBP or anti-V5 antibodies that bind the MBP domain or the V5 tag of the recombinant proteins.

**Figure 1:**
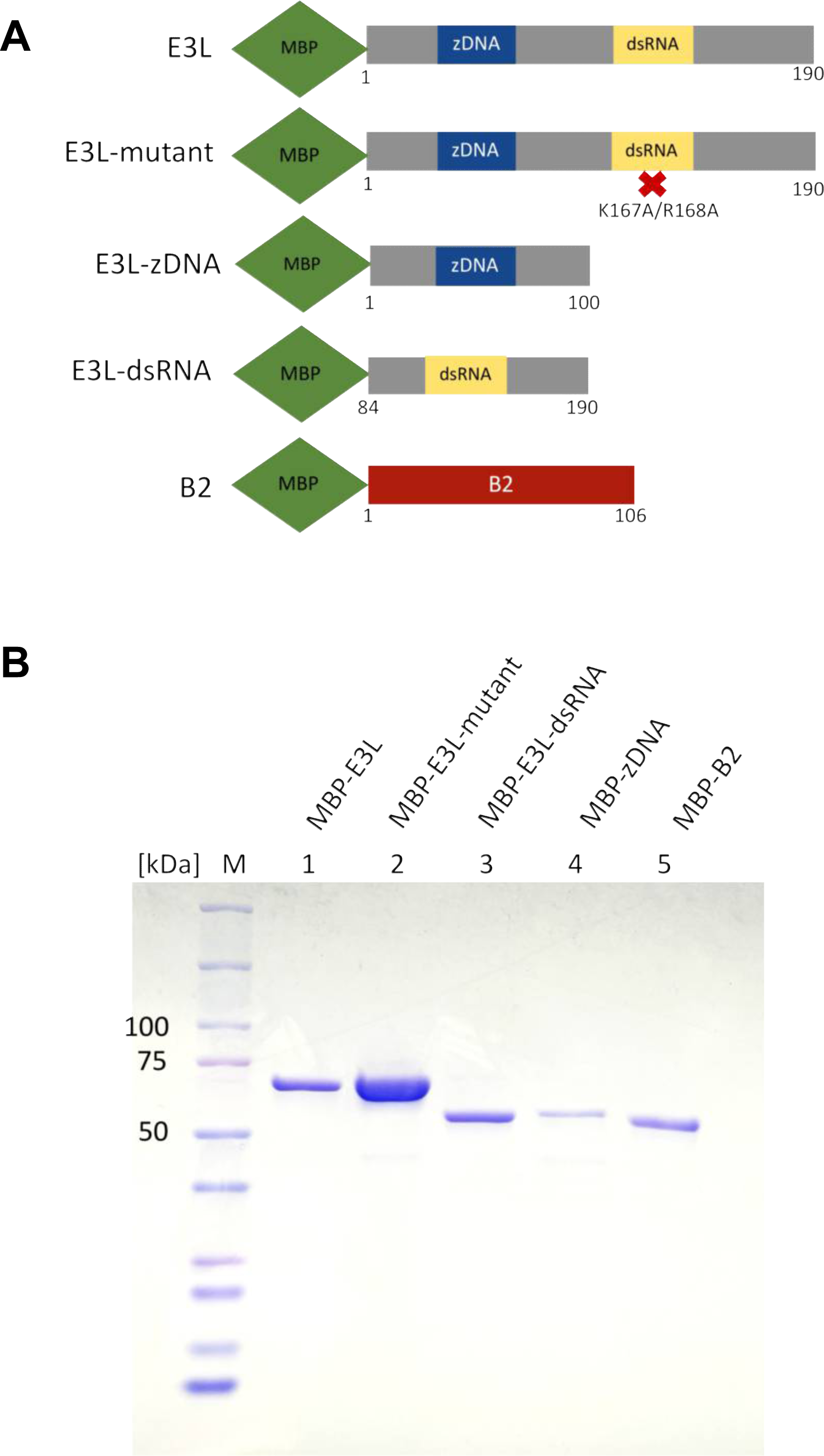
Recombinant Protein production and purification. **(A)** Schematic of the recombinant probes containing various domains of the E3L protein or its derivatives, or B2. All recombinant probes were tagged with the Maltose-binding protein (MBP) and V5 that can be detected using anti-MBV/V5 antibodies. **(B)** Purified recombinant proteins on a polyacrylamide gel electrophoresis stained with Coomassie blue.

### Recombinant E3L protein detects viral dsRNA

The recombinant E3L proteins and known dsRNA markers, J2 and 9D5, were compared for their ability to bind to dsRNAs **(Figure 2)**. To generate dsRNA, human lung-derived epithelial cells (A549 cell line) were infected with Zika virus (ZIKV) from the Flaviviridae family, which is known to generate dsRNA in the cytoplasm during viral replication (16). To visualize the ability of E3L recombinant proteins to detect dsRNA we used mouse anti-MBP or rabbit anti-V5 antibodies and anti-mouse or anti-rabbit antibodies conjugated with Alexa-488. As positive controls, the J2 and 9D5 antibodies were used (17). **Figure 2A** shows that both E3L, E3L-dsRNA and B2 recombinant proteins are detected in Zika infected cells, while E3L-mutant and E3L-zDNA recombinant protein detection was negative. These results suggest that E3L and B2 can detect dsRNA generated during Zika infection **(Figure 2)**. The E3L-mutant K167A-K168A lacking the dsRNA ability was not detected in Zika infected cells suggesting that E3L dsRNA motif is responsible for detecting the signal molecule **(Figure 2A-B)**. The colocalization of E3L with 9D5 and J2 antibodies (**Figure S1**) demonstrate that E3L detects similar dsRNA molecules as J2 and 9D5 (17). Additionally, the higher intensity of signal produced by the E3L and E3L-dsRNA probes compared to J2 and 9D5 antibodies suggests that E3L is a better tool at detecting dsRNA compared to the current standard.

**Figure 2:**
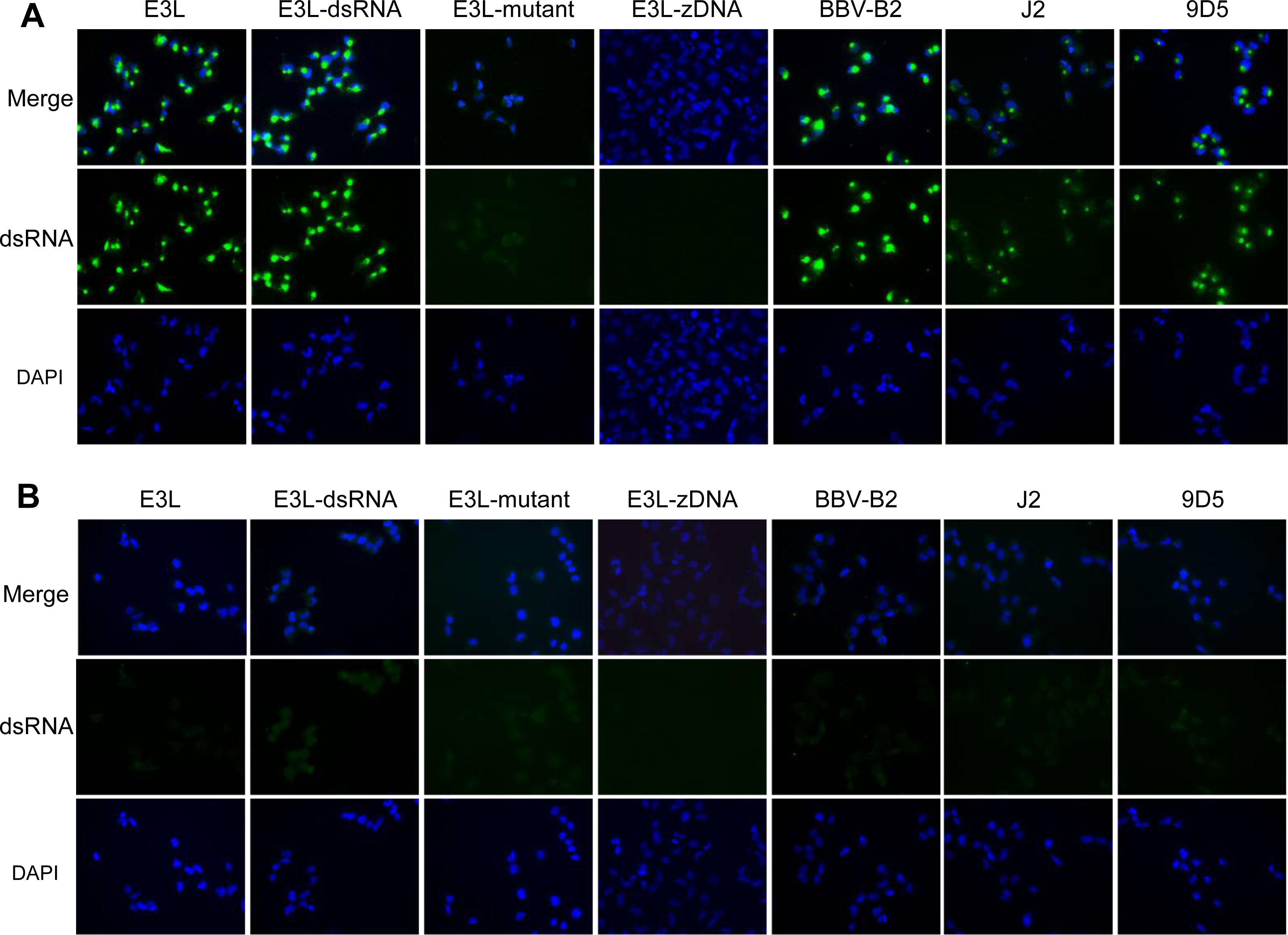
Detection of ZIKV-dsRNA in infected A549 cells. A549 cells were infected with Zika virus (ZIKV) **(A)** or mock **(B)** at a MOI=5 and fixed at 24 hours post-infection (hpi). Samples were processed for immunofluorescence with antibodies or dyes to label nuclei (DAPI, blue) and anti-MPB or V5 antibodies to detect the recombinant proteins (green), J2 (green), or 9D5 (green). Detection of dsRNA by BBV-B2, J2, and 9D5 serve as positive controls.

To characterize whether the recombinant E3L protein binds nonspecifically to all RNAs, or specifically to either ssRNA or dsRNA, we used endoribonucleases RNase I, which degrades ssRNA, and RNase III, which degrades dsRNA **(Figure 3)**. Here, A549 cells infected with ZIKV were fixed by ice-cold methanol and treated with RNase I or RNAse III, followed by immunofluorescence. When we used E3L as the probe, we observed a robust expression of fluorescent signal compared to untreated controls. RNase I treatment did not significantly alter the detection of dsRNA, while RNase III treatment completely abrogated the dsRNA detection. This suggests that the recombinant E3L protein binds specifically to dsRNAs that are generated during virus infections, and not ssRNAs that are present in mammalian cells. As expected, the control 9D5 antibody, which is known to detect dsRNA, also lost expression upon RNase III treatment, further showing that recombinant E3L protein indeed binds to dsRNA and not ssRNA. However, the 9D5 antibody signal was reduced with RNase I treatment, suggesting a lack of specificity to dsRNA. Together, these results show that the recombinant E3L protein can specifically detect dsRNA better than the current commercial marker 9D5.

**Figure 3:**
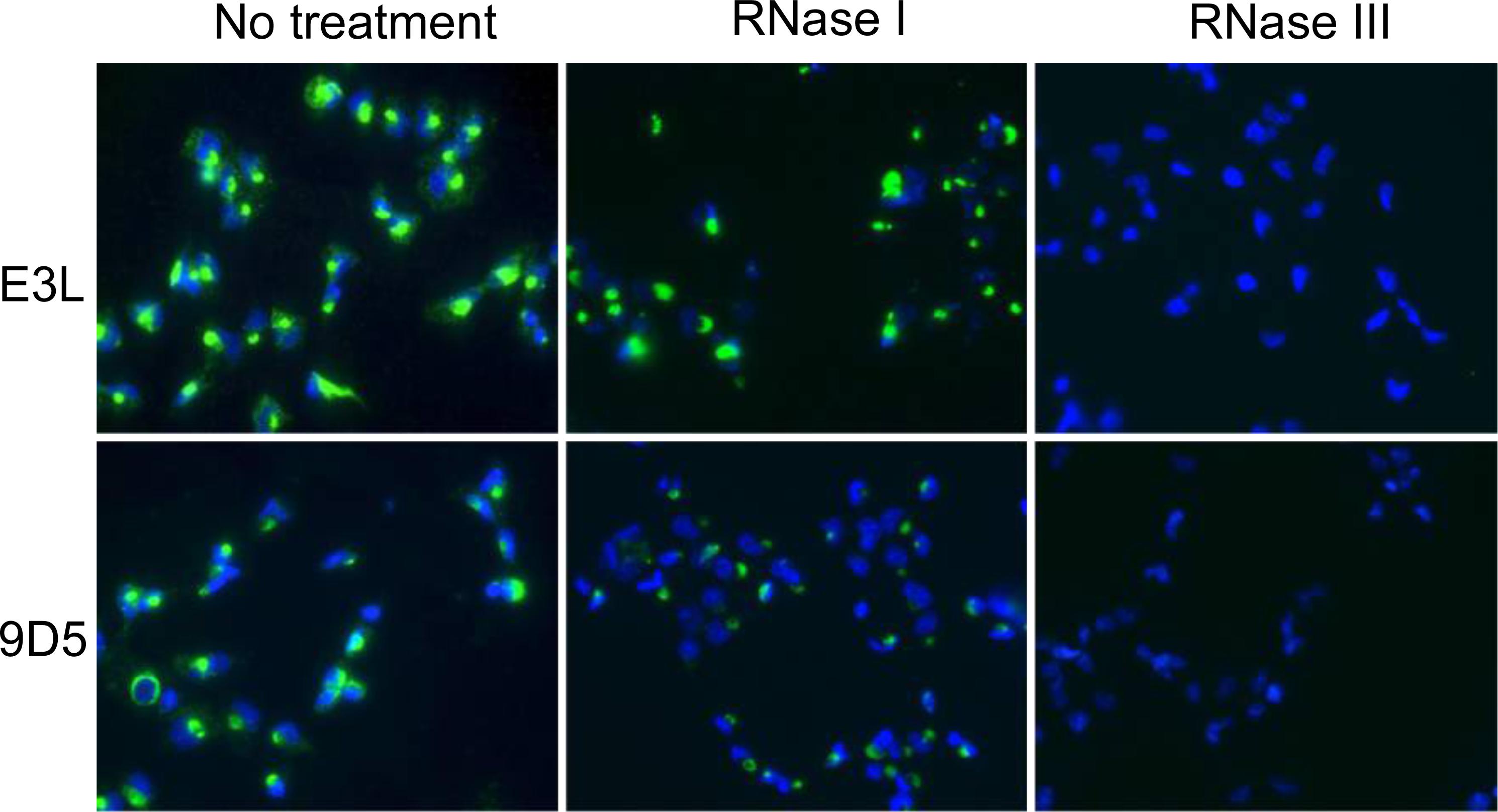
Detection of ZIKV-dsRNA after RNase I/III treatment in A549 cells. A549 cells were infected with ZIKV at MOI=5. At 24 hpi cells were fixed by ice-cold methanol, cells were permeabilized by 0.1% Triton-X100 and then treated with 50U/ml of RNase I or 100U/ml RNase III followed by incubation at 37°C for 2 hours. Samples were labeled with DAPI (blue) and anti-V5/MBP (green) or anti-9D5 (green) antibodies.

### Recombinant E3L protein binds to dsRNA during infections by positive-sense RNA genomes

To determine whether viral dsRNA detection by the recombinant E3L proteins is specific to Zika virus infections, or if it can be used to detect dsRNA from a broader range of viruses, we compared the recombinant E3L protein expression during infections by various viruses **(Figure 4-5)**.

**Figure 4:**
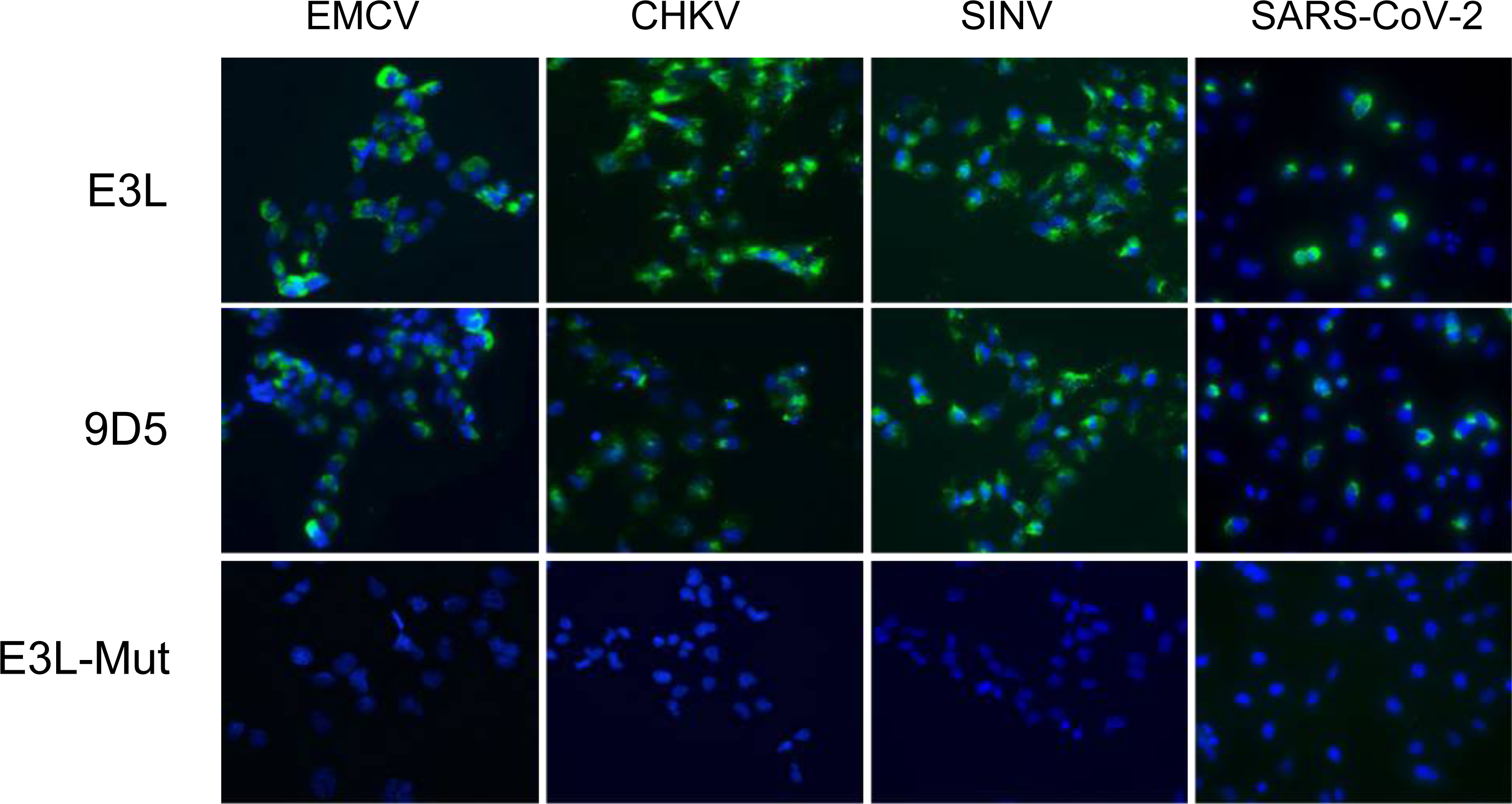
Detection of dsRNA in A549 cells infected with positive-sense RNA viruses. A549^ACE2^ cells were infected with EMCV (MOI=0.1), CHKV, SINV-mCherry, or SARS-CoV-2 viruses at MOI=1 and fixed at 24 hpi. Samples were processed for immunofluorescence with antibodies to label nuclei (DAPI, blue), anti-MBP (green) or anti-V5 (green), or 9D5 (green).

**Figure 5:**
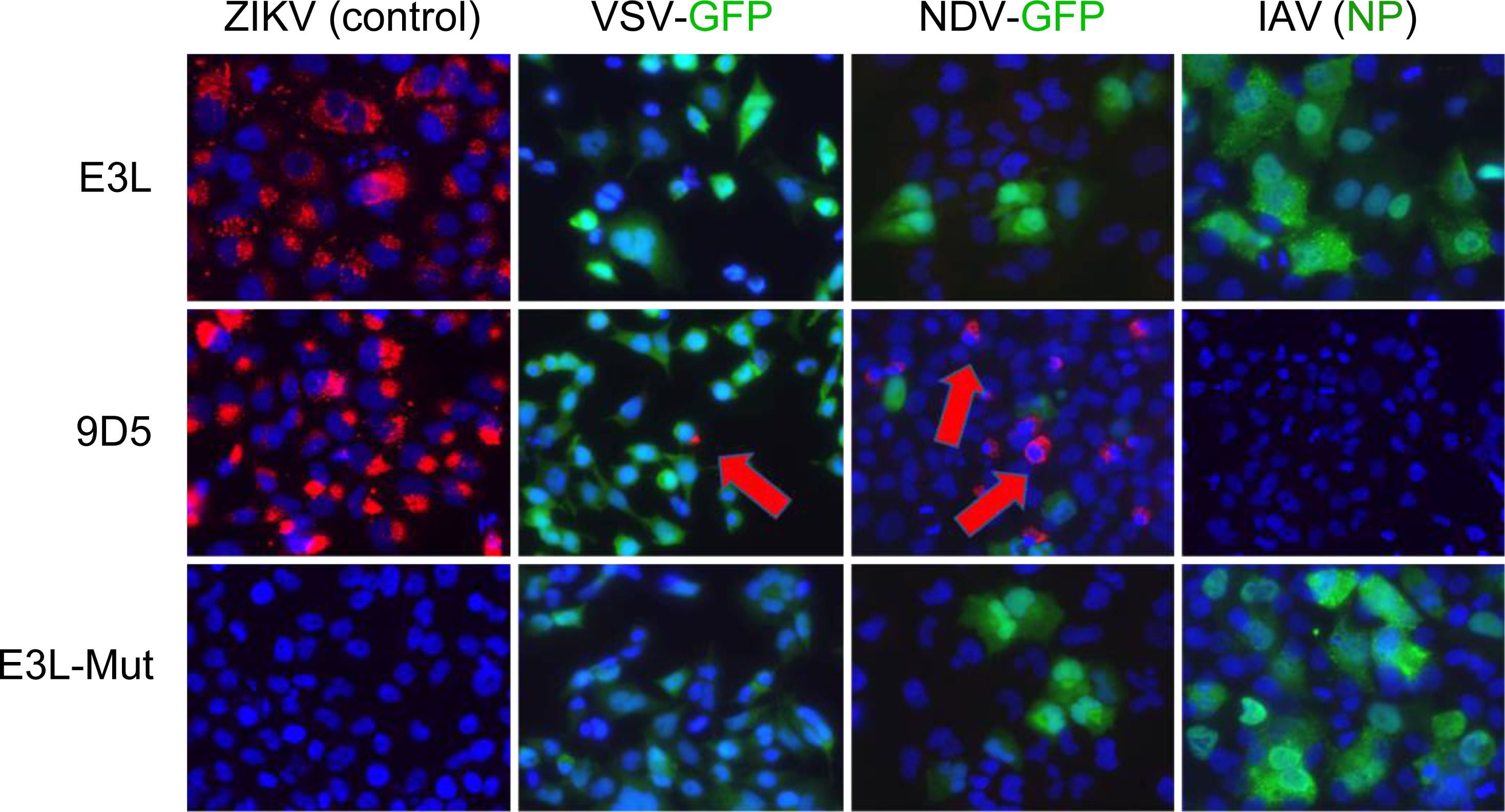
Detection of dsRNA in A549 cells infected with negative-sense RNA viruses. A549 cells were infected with ZIKV (MOI=1, fixed 24 hpi), VSV-GFP (MOI=1, fixed 12 hpi), NDV-GFP (MOI=1, fixed 24 hpi) or IAV (MOI=10, fixed 48 hpi) viruses. Samples were processed for immunofluorescence with DAPI (blue), and anti-V5/anti-MPB/9D5 antibodies (red). The IAV sample was also processed with an anti-NP antibody (green).

To measure the detection of dsRNA generated by positive-sense dsRNA viruses, A549 cells were infected by the following viruses: the encephalomyocarditis virus (EMCV) from the Picornaviridae family, Chikungunya virus (CHKV) and Sindbis virus (SINV) from the Togaviridae family within the Alphavirus genus, and SARS-CoV-2 virus from the Coronaviridae family **(Figure 4)**. While EMCV, CHKV, and SINV infections are susceptible and permissive in A549 cells, SARS-CoV-2 needs the ACE2 receptor that is not abundantly expressed in A549 cells (18). Therefore, A549^ACE2^ cells that overexpress the ACE2 receptor of SARS-CoV-2 were used for infection (19), as the overexpression of ACE2 does not affect infections by EMCV, CHKV, or SINV. All infections were performed at MOI=1 and cells were fixed at 24 hpi. Both the recombinant E3L protein and the 9D5 antibody detected dsRNA generated by all the viruses tested, although the signal was brighter with the recombinant E3L protein compared to the 9D5 antibody. As expected, the recombinant E3L-mutant protein, which lacks the dsRNA binding ability, did not detect dsRNA in any of the infections. This demonstrates that the dsRNA motif of E3L can detect dsRNA from a wide range of viruses composed of (+)ssRNA genomes.

To measure the detection of dsRNA generated by negative-sense viral RNA genomes, we compared infections by recombinant vesicular stomatitis virus (VSV-GFP) from the Rhabdoviridae family, recombinant Newcastle disease virus (NDV-GFP) from the Paramyxoviridae family, and influenza A virus (IAV) from the Orthomyxoviridae family **(Figure 5)**. Upon infection of A549 cells, followed by immunofluorescence assay, we observed that in comparison to Zika-infected cells as a positive control, the E3L recombinant protein was not detected (red color) in cells infected with (-)ssRNA viruses (labeled green with GFP for VSV and NDV, or NP protein for IAV). The 9D5 antibody, however, was present in some infected cells (indicated by red arrows) with VSV-GFP and NDV-GFP but not IAV. As expected, the E3L-mutant probe did not detect dsRNA in the ZIKV control or VSV, NDV, and IAV infections. Together these results demonstrate that the E3L probe specifically detects A-dsRNA from viruses with a positive-sense RNA genome.

### The recombinant E3L-zDNA protein did not recognize Z-NA during IAV infection

Cellular stress from insults such as viral infection can induce the expression of less-familiar genomic structures. For example, infections with IAV can activate innate immune responses that produce Z-RNAs (7). To test whether Z-RNA can be detected by anti-Z-RNA antibodies during infection with viruses other than IAV, we infected A549 cells with viruses in our collection (ZIKV, CHKV, EMCV, SARS-CoV-2, SINV, VSV, NDV) and confirmed that only IAV infection generates Z-RNA **(Figure 6A and B)**. Sensing of Z-RNA however can be prevented by the Z-DNA binding domain of E3L, such as during infection by vaccinia virus (11).

**Figure 6:**
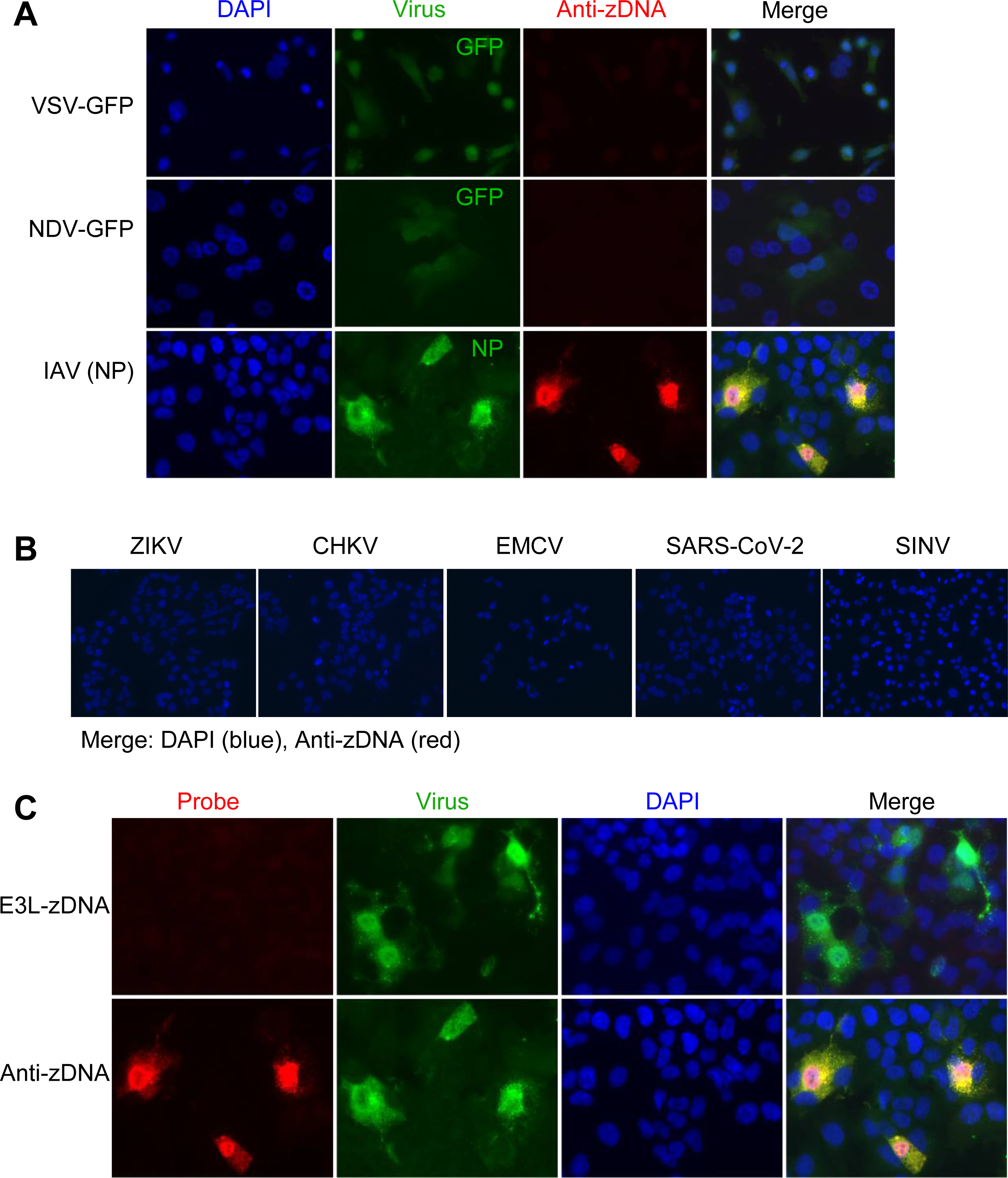
Detection of Z-DNA in A549 cells infected with multiple viruses. **(A)** A549^ACE2^ cells were infected with positive-sense RNA genome viruses: ZIKA, CHKV, SINV, or SARS-CoV-2 viruses at MOI=1 and fixed at 24 hpi. Images are a merge of DAPI (blue) and anti-Z-DNA (red) channels. **(B)** A549 cells were infected with negative-sense RNA genomes VSV-GFP (MOI=1, 12 hpi), NDV-GFP (MOI=1, 24 hpi) or IAV (MOI=10, 48 hpi) viruses. Samples were processed for immunofluorescence with DAPI (blue), anti-V5/MBP antibody (red), or anti-zDNA antibody (red). The green channel is labeled as indicated. **(C)** Detection of zDNA and E3L-zDNA in IAV infected A549 cells. A549 cells were infected with IAV at MOI=10 and fixed at 48 hpi. Samples were fixed for immunofluorescence with DAPI (blue), anti-zDNA (red) antibody, or anti-V5/MBP antibody (green).

To assess whether the recombinant E3L-zDNA protein can recognize the Z-RNA from cells infected with IAV, we infected A549 cells with IAV and performed immunofluorescence with antibody against the nucleocapsid (NP) protein to mark infected cells. The E3L-zDNA recombinant protein did not detect Z-RNA from IAV infection **(Figure 6C)**.

### The recombinant E3L protein binds to endogenous dsRNA in PNPT1 or SUV3L1 KO cells

Regulation of dsRNAs in the mitochondria is mediated by the helicase SUPV3L1 and the endoribonuclease PNPT1 (5). Knockdown of PNPT1 leads to RNA accumulation by inhibiting apoptotic RNA decay (Lui 2018). The RNA helicase SUPV3L1 is part of the RNA degradosome complex and is involved in mitochondrial RNA surveillance and degradation (Paul 2010).

To induce the accumulation of dsRNA in the mitochondria, we used CRISPR/Cas9 to generate PNPT1 or SUPV3L1 knockout A549 cells. We used western blots to confirm efficient PNPT1 protein knockout and partial SUPV3L1 knockout **(Figure 7A)**. The knockout cells were subsequently used to detect dsRNA by the recombinant E3L protein or the 9D5 antibody as a control in the mitochondria (labeled with anti-AIF antibody) by immunofluorescence **(Figure 7B)**. We observed that in wild type A549 cells there was no dsRNA accumulation in the mitochondria, as expected. We found that E3L and 9D5 recognized dsRNA in A549 PTPN1-KO and SUPV3L1-KO cells, suggesting that both the recombinant E3L protein and 9D5 antibody can bind to mitochondrial dsRNA at similar levels.

**Figure 7:**
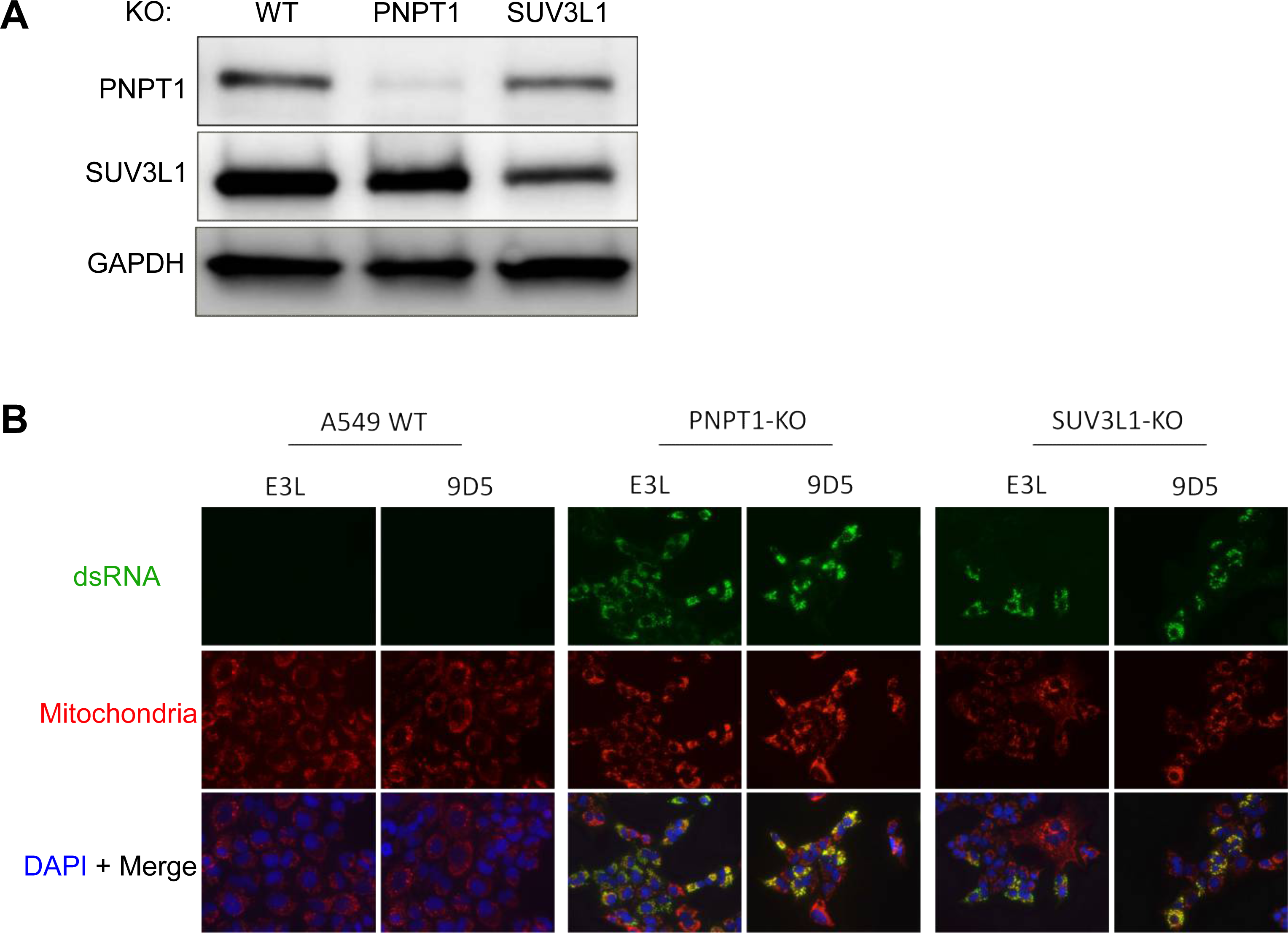
Detection of endogenous mitochondrial dsRNA. **(A)** A549 cells with PNPT1-knockout or SUV3L1-knockout were processed by western blot and stained with PNPT1, SUV3L1, or GAPDH (loading control) antibodies to assay protein knockout. **(B)** The A549 wildtype and knockout cells were fixed and processed for immunofluorescence staining DAPI (blue), 9D5 or anti-V5/MBP antibody (green) and anti-AIF (red) to label mitochondria.

To compare if E3L and 9D5 probes detect similar endogenous dsRNA, we co-stained dsRNA by 9D5 and E3L in SUV3L1 or PNPT1 KO cells. In the SUV3L1-KO cells, the signal from both probes overlapped. In PNPT1-KO cells the signal merged partially, as some cells that were positive for the E3L signal were negative for the 9D5 signal **(Figure S2)**.

### E3L prevents the phosphorylation of PKR as well as cells death in ADAR-p150 KO cells upon interferon treatment

ADAR1 deficiency causes the accumulation of endogenous dsRNA and these dsRNAs are detected by PKR, inducing cell death (20, 21). To assay whether E3L could sequester endogenous dsRNA and prevent sensing of dsRNA by PKR, we transduced ADAR1-p150 KO cells (p150-KO) with lentiviral vectors expressing E3L or E3L-M(K167A/R168A). Upon IFN treatment, phosphorylated PKR (pPKR) was detected in WT and p150-KO cells **(Figure 8A)**. In WT cells, the expression of E3L or E3L-mutant did not affect the level of pPKR **(Figure 8A)**. In p150-KO cells, higher levels of pPKR than in WT cells were detected upon IFN treatment and the level of pPRK was reduced with E3L expression compared to E3L-mutant expression **(Figure 8A)**. These results suggested that E3L could suppress the activation of PKR by endogenous dsRNA through dsRNA binding in p150-KO cells.

**Figure 8:**
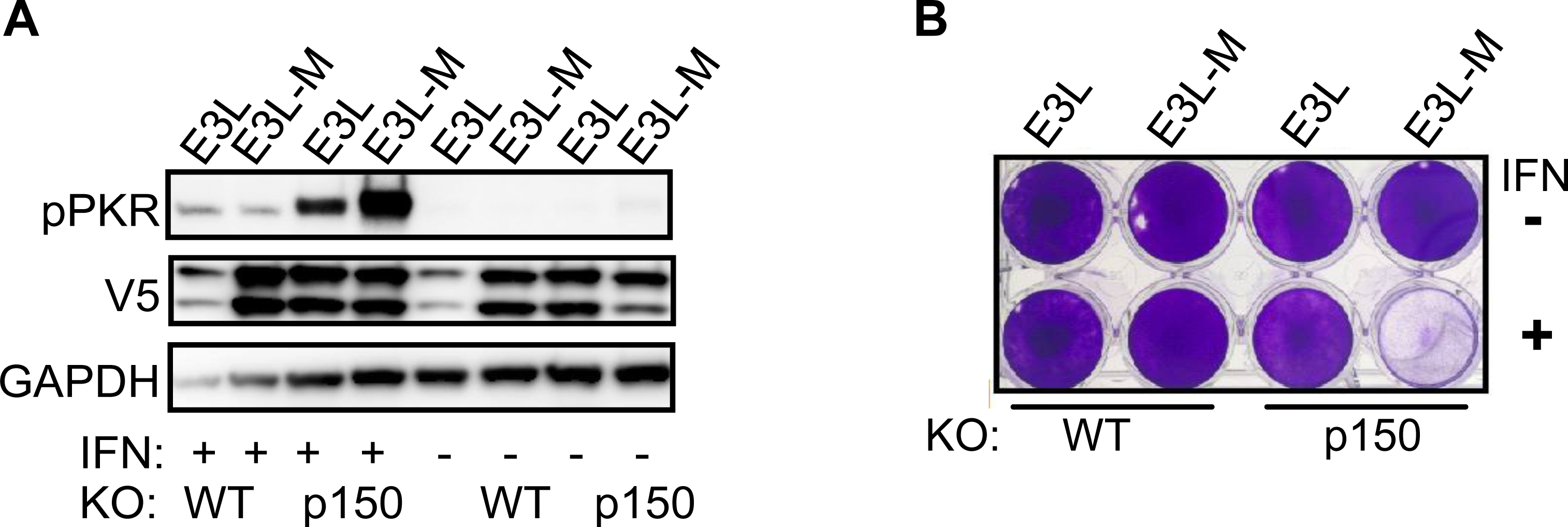
Prevention of endogenous dsRNA-induced cell death by E3L protein. WT or ADAR1-P150-KO cells were transduced with lentiviral vector to express E3L or E3L-M (K167A/R168A). Cells were mock-infected or treated with 1000 U of universal IFN-α for 48 hrs. Cells were harvested for western blotting **(A)**. Cells were fixed by 4% PFA and stained by crystal violet crystal **(B)**.

Activation of PKR could lead to cell death; thus we measured cell death to determine if E3L expression could prevent endogenous dsRNA-induced cell death. Cells were treated with IFN and stained with crystal violet; here, viable cells absorb and stain positive for the violet dye, while dead cells fail to absorb the dye. In WT cells, IFN treatment did not cause cell death **(Figure 8B)**. In p150-KO cells treated with IFN, cells with E3L-mutant expression did not survive while cells with E3L expression survived **(Figure 8B)**.

## Discussion

Detection of dsRNA is crucial for various cellular processes, including the activation of innate immune responses. To date there are some discrepancies in the detection of dsRNA during viral infections. In this study we report the generation and characterization of a novel dsRNA probe, the E3L recombinant protein, which provides significant insight into the biology of RNAs generated during viral infections. First, we have demonstrated that the E3L recombinant protein specifically detects A-dsRNAs, in comparison to the J2 and 9D5 antibodies that nonspecifically detect both dsRNAs and ssRNAs. Moreover, the E3L recombinant protein expression and detection are superior to the commercially available antibodies.

An interesting finding from our studies is that viral infections by positive-sense and negative-sense RNA viruses produce dsRNA with different signatures. The E3L recombinant protein detected dsRNA from (+)ssRNA virus infections but not (-)ssRNA virus infections. While both J2 and 9D5 also detect dsRNAs generated by (+)ssRNA viruses, their specificity is not as strong as E3L recombinant proteins. It is curious to consider the characteristics of (-)ssRNA generated during an infection that evades detection by E3L, J2, and 9D5.

Additionally, during an infection by IAV that is composed of a (-)ssRNA genome, the Z-RNAs generated are not detected by the E3L-zDNA probe, suggesting differences in the types of Z-RNAs generated during infection. It may be that E3L has low affinity to Z-dsRNA, however further experimentation is needed to address the question.

Besides viral dsRNA, we have shown that the E3L recombinant protein also detects mitochondrial dsRNA, suggesting a structural similarity between dsRNA generated by mitochondrial damage and infection by (+)ssRNA genome viruses.

A loss-of-function (LOF) of the highly conserved ADAR family can cause endogenous dsRNA accumulation and MDA5-, OASs- and PKR-mediated pathway activation, and the auto immune disease Aicardi-Goutières syndrome (21–24). To assess the role of E3L in this process we created E3L or E3L-M (lacking dsRNA binding affinity) expressing cells in the background of WT and P150-KO cells. In this system upon INF treatment, E3L-M expressing P150-KO cells died while E3L expressing P150-KO cells survived. This result suggests that E3L could bind endogenous dsRNA and prevent dsRNA from stimulating the innate immune pathways. It suggests that E3L or protein with a similar function could be used to attenuate endogenous dsRNA induced tissue damage.

## Materials and Methods

### Cell lines and viruses

Human A549 or A549^ACE2^ cells were cultured in a Roswell Park Memorial Institute (RPMI) 1640 medium (Gibco catalog no. 11875) supplemented with 10% (vol/vol) FBS, 100 U/mL penicillin, and 100 μg/mL streptomycin. The construction of A549^ACE2^ cell line was described in a previous study (19).

Zika virus isolates MR766 (GenBank accession no. KX377335) was obtained from Robert Tesh, University of Texas Medical Branch, Galveston, TX, and propagated in Vero cells. The encephalomyocarditis virus (EMCV) (25) was obtained from Dr. Ian M. Kerr (Cancer Research United, Kingdom, London, United Kingdom); NDV-GFP was obtained from Dr. Luis Martinez-Sobrido (Texas Biomedical Research Institute, San Antonio, TX, USA) The IAV H1N1 strain PR8 was obtained from Dr. Scott Hensley (University of Pennsylvania, Philadelphia, PA, USA) and grown in chicken eggs as described previously (26). Girdwood G100 (SINV), was obtained from Dr. Mark Heise (University of North Carolina, Chapel Hill) and was prepared in BHK cells as previously described (27). SARS-CoV-2 (USA-WA1/2020 strain) was obtained from the Centers for Disease Control and Prevention and obtained through BEI Resources, NIAID, NIH: SARS-related coronavirus 2, 5 Isolate USA-WA1/2020, NR-52281 and propagated in Vero-E6 cells. The genome RNA sequenced was found to be identical to GenBank: MN985325.1. Infections and manipulations of SARS-CoV-2 were conducted in a biosafety level 3 (BSL-3) laboratory using appropriate and approved personal protective equipment and protocols. CHKV was obtained from Dr. Sara Cherry (University of Pennsylvania, Philadelphia, PA, USA).

### Construction of plasmids and pseudolentivirus

The cDNA of E3L of vaccinia was obtained from Dr. Bernard Moss’ lab at the NIH. The cDNA of Black Beetle virus B2 with V5 tag (black beetle virus) was synthesized by GeneArt. Primers were listed in Table S1. The E3L-Mutant (K167A R168A) was generated by fusion PCR. Briefly, primers pMal-E3L-For and M-rev (bearing K167A R168A mutation) were used to amplify the E3L-M-F1, primers M-for (bearing K167A R168A mutation) and pMal-V5-Rev were used to amplify E3L-M-F2. The F1 and F2 (with overlap sequences) were used as a template for fusion PCR, and primers pMal-E3L-For and pMal-V5-Rev were mixed with F1 and F2 also. For cloning of the target DNA into the pMal-C2 vector, primers pMal-E3L-For and pMal-V5-Rev were used to amplify E3L, and primers pMal-E3L-For and zDNA-Rev were used to amplify E3L-zDNA, primers dsRNA-For and pMal-V5-Rev were used amplify E3L-dsRNA, primers B2-For and pMal-V5-Rev were used to amply B2. All the DNA fragments were cloned into pMal-C2 vector between SalI and XbaI restriction sites. The plasmids were named pMal-E3L, pMal-E3L-mutant pMal-E3L-zDNA, pMal-E3L-dsRNA, pMal-B2. E3L and E3L-mutant were amplified by primers pLenti-E3L-for and pLenti-E3L-rev and were cloned into pLenti between XbaI and SalI restriction site. The plasmids were named pLenti-E3L, pLenti-E3L-M.

The oligonucleotide sequences to be used for the generation of small guide RNAs (sgRNAs) for KOs SUV3L1 and PNPT1genes are shown in Table S2. A pair of forward and reverse oligonucleotides for the generation of each sgRNA (synthesized by IDT) were annealed by published methods (28) and were inserted into pLenti-CRISPR (Addgene) between BsmBI restriction sites. The resulting plasmids are named pLenti-sgSUV3L1 (targeting the SUV3L1 gene), pLenti-sgPNPT1 (targeting the PNPT1 gene).

For packaging of pseudolentiviruses, 1□×□10^6^ HEK 293T cells were plated in one well of a 6-well plate, and the next day transfected with 5□μg pLenti or pLenti-CRISPR (with sgRNA), 3.5□μg psPAX2, and 1.25□μg of pCMV-VSV-G (obtained from Paul Bates, University of Pennsylvania) using Lipofectamine 2000 (Invitrogen) (24□μl in 250□μl of DMEM). The supernatants were harvested at 24 and 48 h post-transfection and stored at −80°C, and the 48-h supernatants were used for further KO experiments.

### Generation of PNPT1 or SUV3L1 KO cells by CRISPR/Cas9 techniques

The methods for the construction of single gene knockout (KO) A549 cells using Lenti-CRISPR were described previously (29).

### Recombinant protein production and purification

pMal-E3L, pMal-E3L-mutant pMal-E3L-zDNA, pMal-E3L-dsRNA, pMal-B2 were transformed into NEB-Express competent Escherichia coli, and IPTG was added to the bacterial cultures to induce expression of the proteins. First, the E. coli was harvested by centrifuging at 6000 rpm for 15 min at 4 °C. The pellets were collected, and 4 ml of lysis buffer was added. The mixtures were first incubated in a 37 °C water bath for 30 min and then incubated in a Dry ice-70% ethanol bath for 5-10 min. To lyse the cells completely, the lysates were thawed in cold water, treated with 8ul of 1M MgCl_2_ and 8ul of DNase I, and vortexed until the solution was no longer viscous. By spinning down at 20000xg at 4C for 20 min, supernatants containing recombinant proteins were harvested. The recombinant E3L proteins and their derivatives were purified from the supernatants by washing through a 5ml MBPTrap^TM^ HP column (from Cytiva).

### Polyacrylamide gel electrophoresis

Prior to SDS-polyacrylamide gel electrophoresis (SDS-PAGE), the samples were mixed with SDS-PAGE protein loading dye 4x (Invitrogen) and boiled at 95°C for 5 min. The protein samples were separated in 4 −15% acrylamide gels by SDS-PAGE using a Bio-Rad Mini-Protean Tera cell system (Bio-Rad). At the same time, 5 μL of pre-stained SDS-PAGE Standards (Bio-Rad) were loaded in each gel run. Electrophoresis was performed at room temperature for approximately 1 hour using a constant voltage (100V) in a 1X solution of an SDS running buffer (Invitrogen) until the dye almost reached the bottom of the gel. The gel carrying the recombinant proteins and their derivatives was eventually stained with Coomassie Blue (Bio-Rad) for 1 hour and was de-stained with a de-staining solution (20% methanol and 10% acetic acid in water).

### Western immunoblotting

Cells were mock treated or treated with 1000U of universal IFN-IZl (PBL Assay Science Cat # 11200-2) Cells were washed twice with ice-cold PBS and lysates harvested with lysis buffer (1% NP-40, 2 mM EDTA, 10% glycerol, 150 mM NaCl, 50 mM Tris HCl) supplemented with protease inhibitors (Roche cOmplete mini-EDTA-free protease inhibitor) and phosphatase inhibitors (Roche PhosStop Easy Pack). After 5 min, lysates were harvested, incubated on ice for 20 min, and centrifuged for 20 min at 4°C, and supernatants were mixed 3:1 with 4x Laemmli sample buffer. Samples were heated at 95°C for 5 min and then separated by 4 to 15% SDS-PAGE and transferred to polyvinylidene difluoride (PVDF) membranes. Blots were blocked with 5% BSA and probed with antibodies (table below) diluted in the same block buffer. Primary antibodies were incubated overnight at 4C or for 1 hour at room temperature. Blots were visualized using Thermo Scientific SuperSignal west chemiluminescent substrates (Cat #: 34095 or 34080). Blots were probed sequentially with antibodies and in between antibody treatments stripped using Thermo Scientific Restore western blot stripping buffer (Cat #: 21059).

PNPT1 and SUV3L1 proteins were detected using anti-PNPT1 antibodies and anti-SUV3L1 antibodies at 1:1000 (both obtained from Abcam, Cat #: ab176845 and Cat #: ab157109). pPKR, V5 tagged proteins, and GAPDH were detected using anti-pPKR (Abcam, Cat#32036), anti-V5(Cell Signaling Technology, Cat#13202S), and anti-GAPDH antibodies (Cell Signaling Technology, Cat#2118) at 1:1000. Secondary antibodies were HRP linked goat anti-rabbit IgG (Cell Signaling Technology 7074S) diluted in 5% milk/TBST blocking buffer at 1:3000.

### Viral infection and immunofluorescent staining

Wildtype A549 cells were cultured in an RPMI 1640 medium with 10% FBS on 8-well chamber slides (Millicell) on Day 0. On Day 1, the cells were incubated with viruses which were diluted in a 2% FBS RPMI 1640 medium. At indicated times post-infection, the cells were fixed with 4% paraformaldehyde (PFA) for 20 min at room temperature. The cells were then permeabilized with 0.1% Triton X-100 in PBS for 10 min and blocked with 5% bovine serum albumin (BSA) in PBS for 10 min at room temperature. The recombinant proteins (1ug/ml) or antibodies J2 (Scions, 1:1000), 9D5 (EMD Millipore, Cat# 3361, 1:10) for dsRNA or goat anti zNA polyclonal antibodies (Novus, NB100-749, 1:100) or anti-AIF (Cell Signal, Cat# 5318, 1:500) mitochondria maker, or anti-influenza A virus NP antibody (Abcam Cat# ab128193) were then diluted and added into each well to bind target molecules in the cells. After incubating for 1 hour and washing three times with PBS, the primary antibodies were diluted in a 2% BSA blocking buffer and incubated on a rocker at room temperature for 1 hour. The cells were washed three times with a PBS buffer 1X and then incubated with mouse anti-MBP (NEB, Cat# E8032L) or rabbit anti-V5 (Cell Signaling Technology, Cat#13202S) (for the recombinant proteins). The cells were washed 3 times and then were incubated with goat anti-mouse antibodies, goat anti-rabbit antibodies, donkey anti-sheep antibodies, conjugated with Alexa-488 or Alexa-594, (Life Technology, Cat#s A11029, A11020, A 11034, A11037, A11015, A11016) diluted (1:800) in blocking buffer at room temperature for 1 hour with rocking. The cells were finally washed three times with PBS, and the nucleus was stained with DAPI (4=,6-diamidino-2-phenylindole) diluted in PBS (1:3000). Thin glass coverslips were mounted onto the cell slides for analysis by confocal microscopy, and to preserve the fluorescent signals, a drop of an anti-fluorescence quenching agent was added between each coverslip and cell slide.

Recombinant E3L protein and its derivatives were detected using anti-MBP antibodies at 1:800 or anti-V5 rabbit serum at 1:500 (both obtained from Bio-Rad, catalog numbers Ab1513 and Ab1266). An anti-influenza A N-protein antibody (Abcam, Cat# ab128193) was used to detect the influenza A virus. Secondary antibodies were all highly cross-adsorbed IgG from Invitrogen: goat anti-rabbit AF594 (Cat# AA12567), goat anti-rabbit AF488 (Cat# AA11045), goat anti-mouse AF594 (Cat# A12839), goat anti-mouse AF488 (Cat# A11452), donkey anti-sheep AF594 (Cat# A11016), and donkey anti-sheep AF488 (Cat# A11015).

### RNase treatment

For the RNase treated cells, fixed infected A549 cells with ice-cold methanol at −20C for 15 min. Washed cells with cold PBS three times. After permeabilization with 0.1% Triton X100 for 10 min at room temperature, washed with DPBS three more times. First, diluted RNase I (stock 100 U/ul, Ambion, Cat# AM2295) in PBS to 50 U/ml, and diluted RNase III (stock I U/ul, Ambion Cat# AM2290) to 100 U/ml with buffer and ddH2O (with a volume ratio of 1:1:8). Then, added 200ul of diluted RNase to each well and incubated at 37C for 2hr. The following steps are the same as that of immunofluorescent staining.

### Crystal violet cell death assay

Cells were transduced by Lenti-E3L or Lenti-E3L-M at 48 hrs post transducing, cells were mock treated or treated with 100U of universal IFN-IZl. At 72 hr post-treatment, cells were fixed with 4% paraformaldehyde and stained with 1% crystal violet.

## Acknowledgements

This work was supported by National Institutes of Health grants R01 AI104887 (SRW) and NST was supported in part by T32NS43126-18.

---

**Figure S1:**
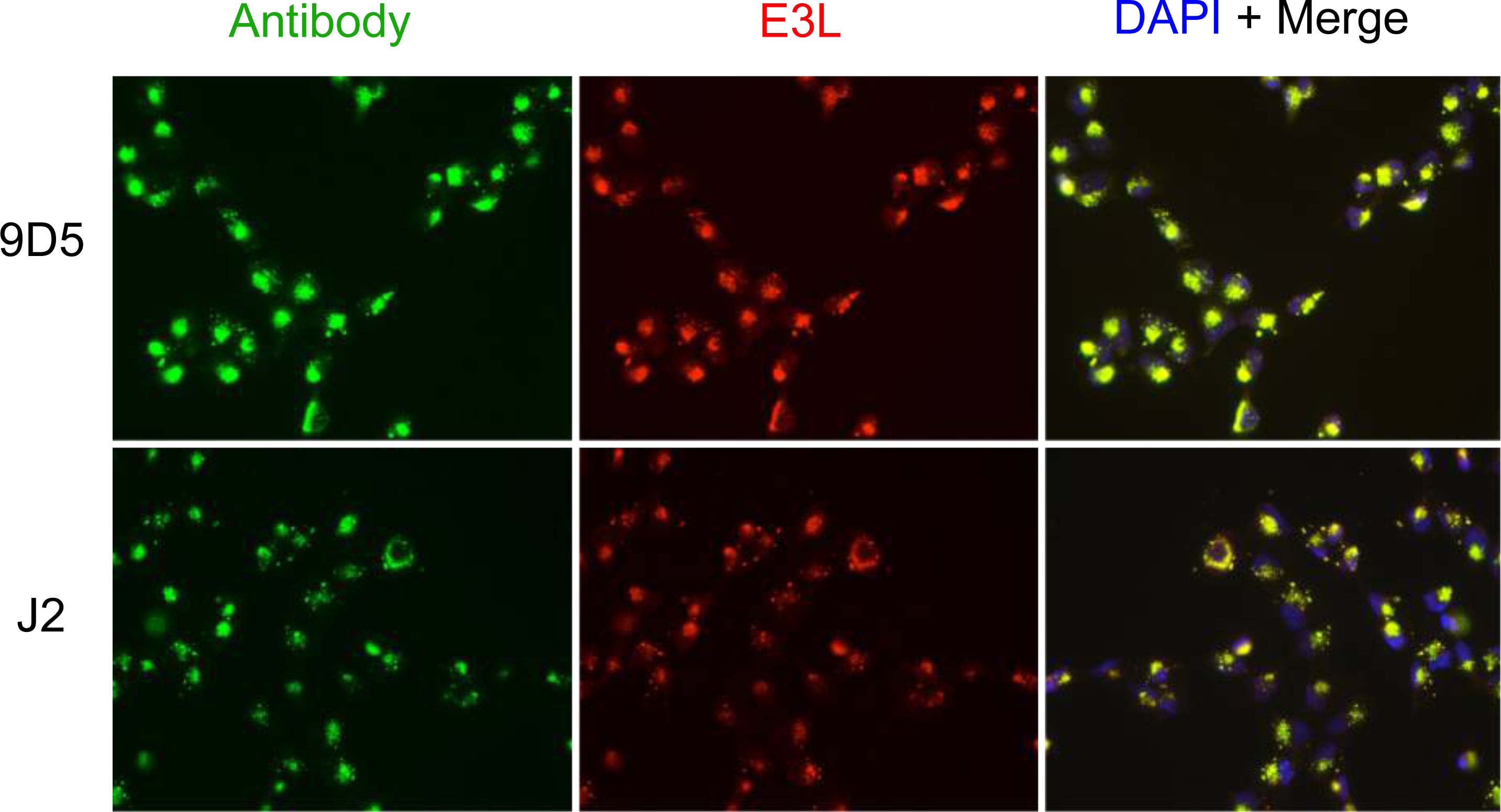
Co-localization of E3L with 9D5 and J2 in A549 cells infected with ZIKV. A549 cells were infected with ZIKV at MOI=5 and fixed at 24 hpi. Samples were processed for immunofluorescence with DAPI (blue), anti-V5/MBP antibody (red), and anti-9D5 or anti-J2 antibodies (green).

**Figure S2:**
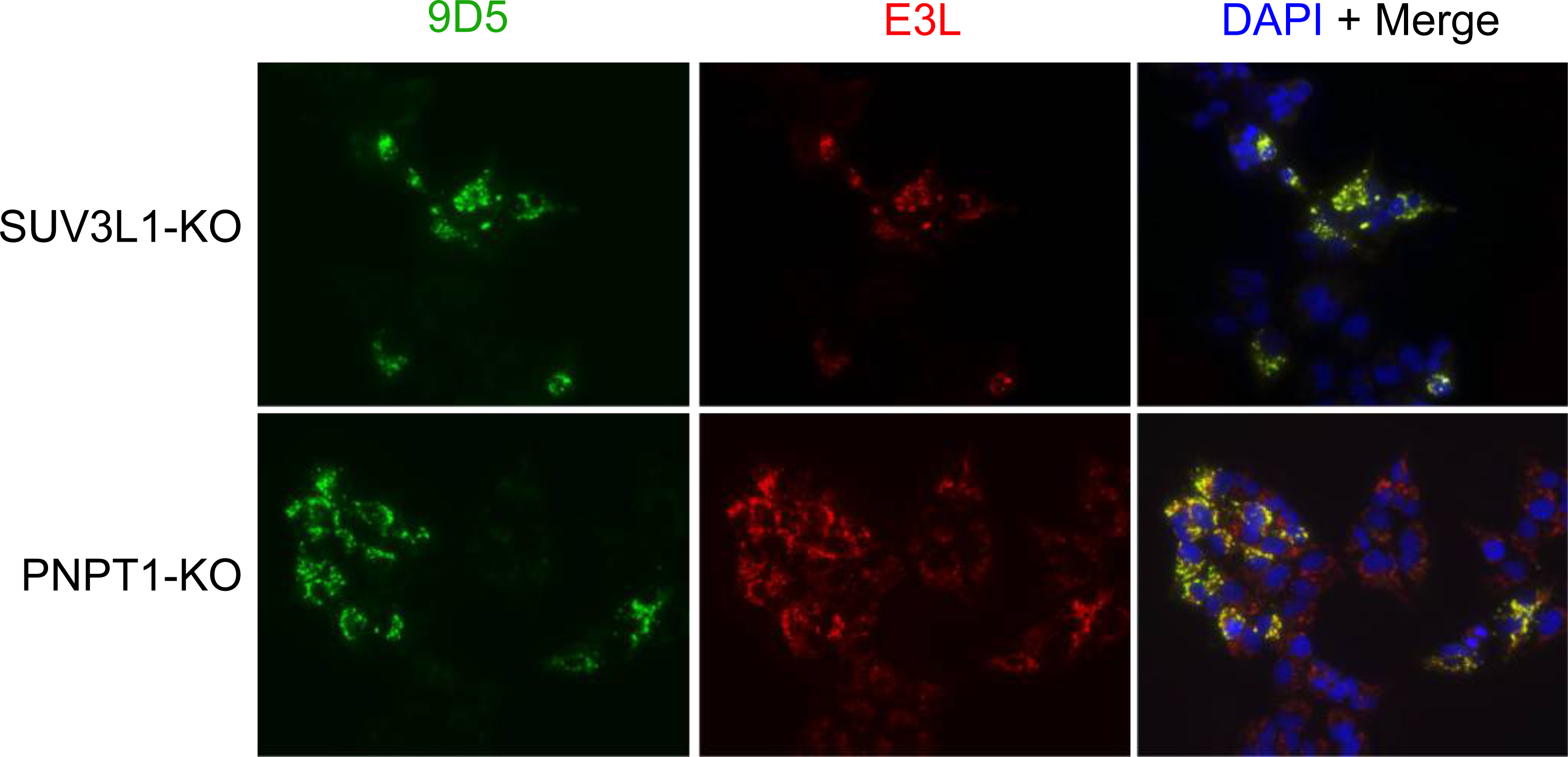
Co-localization of E3L with 9D5 and J2 in A549 SUV3L1- and PNPT1-knockout cells. A549 cells with SUV3L1 or PNPT1-knockouts were fixed and processed for immunofluorescence with DAPI (blue), anti-V5/MBP (red), or anti-9D5 (green) antibodies.

**Table 1.**
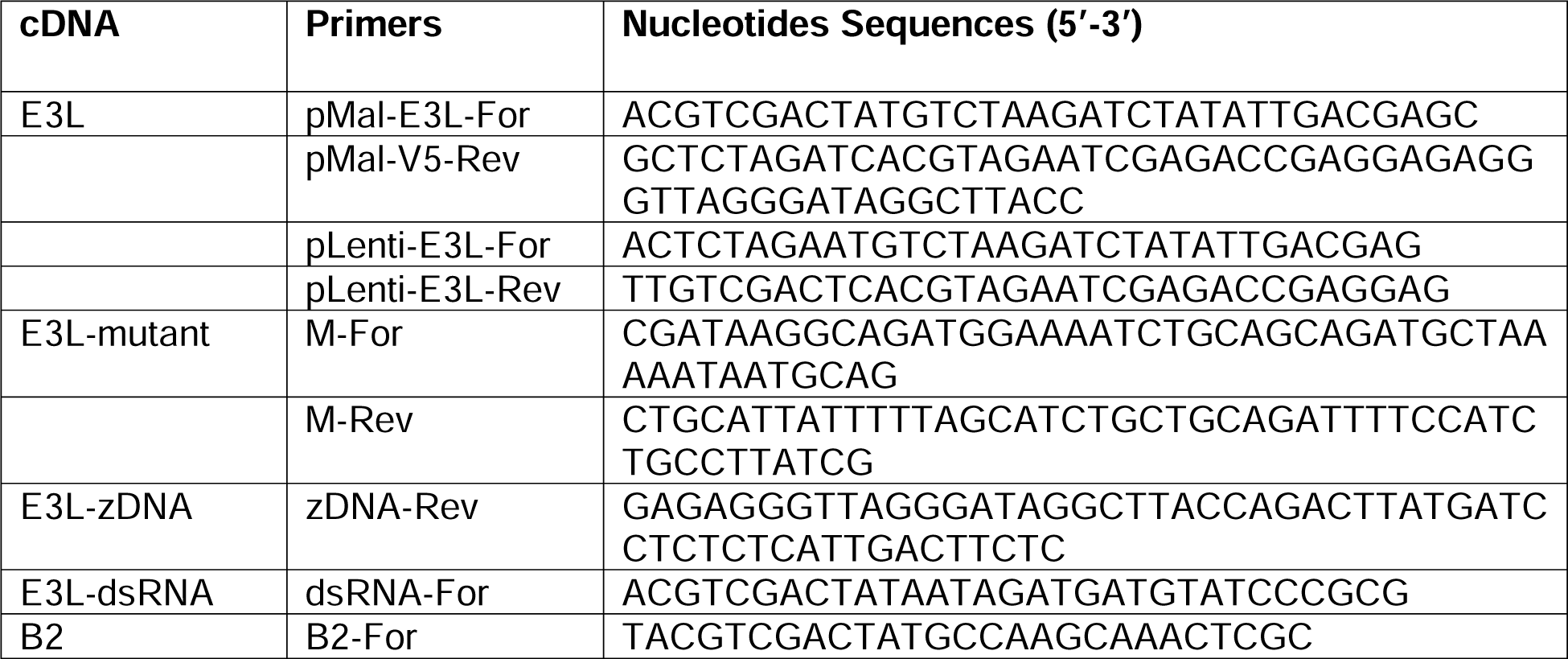
Primers for cloning the cDNA of E3L, E3L-mutant, E3L-zDNA, E3L-dsRNA, B2 to pMal-C2 and pLenti.

**Table 2.**
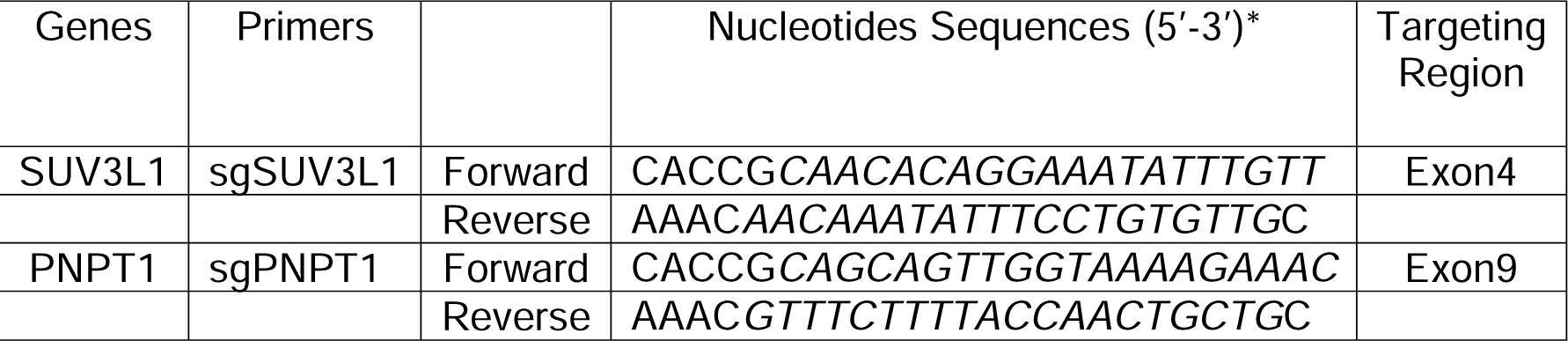
Construction of the plasmids for knockout human SUV3L1 and PNPT1 by CRISPR/Cas9.

## References

1. Monsion B, Incarbone M, Hleibieh K, Poignavent V, Ghannam A, Dunoyer P, et al. Efficient Detection of Long dsRNA in Vitro and in Vivo Using the dsRNA Binding Domain from FHV B2 Protein. Frontiers in plant science. 2018;9:70.

2. Gantier MP, Williams BR. The response of mammalian cells to double-stranded RNA. Cytokine Growth Factor Rev. 2007;18(5-6):363–71.

3. Kuriakose T, Kanneganti TD. ZBP1: Innate Sensor Regulating Cell Death and Inflammation. Trends in immunology. 2018;39(2):123–34.

4. Jiao H, Wachsmuth L, Kumari S, Schwarzer R, Lin J, Eren RO, et al. Z-nucleic-acid sensing triggers ZBP1-dependent necroptosis and inflammation. Nature. 2020;580(7803):391-5.

5. Dhir A, Dhir S, Borowski LS, Jimenez L, Teitell M, Rötig A, et al. Mitochondrial double-stranded RNA triggers antiviral signalling in humans. Nature. 2018;560(7717):238-42.

6. Herrero-Galán E, Fuentes-Perez ME, Carrasco C, Valpuesta JM, Carrascosa JL, Moreno-Herrero F, et al. Mechanical Identities of RNA and DNA Double Helices Unveiled at the Single-Molecule Level. Journal of the American Chemical Society. 2013;135(1):122–31.

7. Zhang T, Yin C, Boyd DF, Quarato G, Ingram JP, Shubina M, et al. Influenza Virus Z-RNAs Induce ZBP1-Mediated Necroptosis. Cell. 2020;180(6):1115–29.e13.

8. Herbert A. Z-DNA and Z-RNA in human disease. Communications Biology. 2019;2(1):7.

9. Jiao H, Wachsmuth L, Wolf S, Lohmann J, Nagata M, Kaya GG, et al. ADAR1 averts fatal type I interferon induction by ZBP1. Nature. 2022;607(7920):776-83.

10. Koehler H, Cotsmire S, Langland J, Kibler KV, Kalman D, Upton JW, et al. Inhibition of DAI-dependent necroptosis by the Z-DNA binding domain of the vaccinia virus innate immune evasion protein, E3. Proceedings of the National Academy of Sciences of the United States of America. 2017;114(43):11506-11.

11. Koehler H, Cotsmire S, Zhang T, Balachandran S, Upton JW, Langland J, et al. Vaccinia virus E3 prevents sensing of Z-RNA to block ZBP1-dependent necroptosis. Cell host & microbe. 2021;29(8):1266–76.e5.

12. Marq JB, Hausmann S, Luban J, Kolakofsky D, Garcin D. The double-stranded RNA binding domain of the vaccinia virus E3L protein inhibits both RNA- and DNA-induced activation of interferon beta. The Journal of biological chemistry. 2009;284(38):25471–8.

13. Richardson SJ, Willcox A, Hilton DA, Tauriainen S, Hyoty H, Bone AJ, et al. Use of antisera directed against dsRNA to detect viral infections in formalin-fixed paraffin-embedded tissue. J Clin Virol. 2010;49(3):180–5.

14. Mateer E, Paessler S, Huang C. Confocal Imaging of Double-Stranded RNA and Pattern Recognition Receptors in Negative-Sense RNA Virus Infection. Journal of visualized experiments: JoVE. 2019(143).

15. Qi N, Cai D, Qiu Y, Xie J, Wang Z, Si J, et al. RNA Binding by a Novel Helical Fold of B2 Protein from Wuhan Nodavirus Mediates the Suppression of RNA Interference and Promotes B2 Dimerization. Journal of virology. 2011;85(18):9543–54.

16. Whelan JN, Li Y, Silverman RH, Weiss SR. Zika Virus Production Is Resistant to RNase L Antiviral Activity. Journal of virology. 2019;93(16):e00313–19.

17. Mateer EJ, Paessler S, Huang C. Visualization of Double-Stranded RNA Colocalizing With Pattern Recognition Receptors in Arenavirus Infected Cells. Front Cell Infect Microbiol. 2018;8:251.

18. Zhou P, Yang X-L, Wang X-G, Hu B, Zhang L, Zhang W, et al. A pneumonia outbreak associated with a new coronavirus of probable bat origin. Nature. 2020;579(7798):270-3.

19. Li Y, Renner DM, Comar CE, Whelan JN, Reyes HM, Cardenas-Diaz FL, et al. SARS-CoV-2 induces double-stranded RNA-mediated innate immune responses in respiratory epithelial-derived cells and cardiomyocytes. Proceedings of the National Academy of Sciences. 2021;118(16):e2022643118.

20. Pfaller CK, Li Z, George CX, Samuel CE. Protein kinase PKR and RNA adenosine deaminase ADAR1: new roles for old players as modulators of the interferon response. Current opinion in immunology. 2011;23(5):573–82.

21. Chung H, Calis JJA, Wu X, Sun T, Yu Y, Sarbanes SL, et al. Human ADAR1 Prevents Endogenous RNA from Triggering Translational Shutdown. Cell. 2018;172(4):811–24.e14.

22. Rice GI, Kasher PR, Forte GM, Mannion NM, Greenwood SM, Szynkiewicz M, et al. Mutations in ADAR1 cause Aicardi-Goutières syndrome associated with a type I interferon signature. Nat Genet. 2012;44(11):1243–8.

23. Li Y, Banerjee S, Goldstein SA, Dong B, Gaughan C, Rath S, et al. Ribonuclease L mediates the cell-lethal phenotype of double-stranded RNA editing enzyme ADAR1 deficiency in a human cell line. Elife. 2017;6.

24. Liddicoat BJ, Piskol R, Chalk AM, Ramaswami G, Higuchi M, Hartner JC, et al. RNA editing by ADAR1 prevents MDA5 sensing of endogenous dsRNA as nonself. Science. 2015;349(6252):1115-20.

25. Hoskins JM, Sanders FK. Propagation of mouse encephalomyocarditis virus in ascites tumour cells maintained in vitro. Br J Exp Pathol. 1957;38(3):268–72.

26. Cooper DA, Banerjee S, Chakrabarti A, Garcia-Sastre A, Hesselberth JR, Silverman RH, et al. RNase L targets distinct sites in influenza A virus RNAs. Journal of virology. 2015;89(5):2764–76.

27. Suthar MS, Shabman R, Madric K, Lambeth C, Heise MT. Identification of Adult Mouse Neurovirulence Determinants of the Sindbis Virus Strain AR86. Journal of virology. 2005;79(7):4219–28.

28. Ran FA, Hsu PD, Wright J, Agarwala V, Scott DA, Zhang F. Genome engineering using the CRISPR-Cas9 system. Nature protocols. 2013;8(11):2281–308.

29. Li Y, Banerjee S, Wang Y, Goldstein SA, Dong B, Gaughan C, et al. Activation of RNase L is dependent on OAS3 expression during infection with diverse human viruses. Proceedings of the National Academy of Sciences. 2016;113(8):2241–6.

